# Calcium- and Phosphorus-Supplemented Diet Increases Bone Volume After Four Weeks of High-Speed Treadmill Exercise in Adult Mice

**DOI:** 10.1101/2020.10.13.337873

**Authors:** Michael A. Friedman, David H. Kohn

## Abstract

Exercise has long-lasting benefits to bone health. Bone mass and the ability to exercise both decline with age, making it ideal to exercise earlier in life and to maximize gains in bone mass. Increasing strain on bone and frequency of loading during exercise can increase bone formation rate and cross-sectional area. Combining a short-term exercise program with a calcium- and phosphorus-supplemented diet increases cortical bone tissue mineral content (TMC) and area more than exercise alone in adult mice. It was hypothesized that combining high-speed running with a mineral-supplemented diet would lead to greater cortical TMC and area than high-speed running on a standard diet and low-speed running on a supplemented diet after 4 weeks. Male, 15-week old mice were assigned to 7 groups – a baseline group, non-exercised groups fed a control or supplemented diet, low-speed exercised groups fed a control or supplemented diet, and high-speed exercised groups fed a control or supplemented diet. Exercise consisted of 4 weeks of daily treadmill running for 20 min/day at 12 m/min or 20 m/min for low- and high-speed exercise, respectively. High-speed exercised mice had significantly lower body weight and lower tibial length after 4 weeks. Cortical TMC and area were significantly higher in high-speed exercised mice on the supplemented diet than high-speed exercised mice on the control diet. Trabecular bone volume (BV) and bone density were significantly higher in all groups on the supplemented diet than groups on the control diet, regardless of exercise. For mice on the control diet, non-exercised mice had significantly lower trabecular BV than baseline, while both speeds of exercise prevented this decline. There were few effects of exercise or diet on mechanical properties. For mice on the control diet, exercise significantly decreased serum PINP/CTX ratio on day 9 which may be preventing exercise from increasing bone mass or strength after only 4 weeks. For non-exercised mice on the supplemented diet, the serum PINP/CTX ratio on day 30 was significantly greater than for exercised mice, suggesting the supplemented diet may also lead to significantly greater bone mass in non-exercised mice if these interventions were extended beyond 4 weeks. Increasing treadmill speed can lower body weight while maintaining cortical and trabecular bone mass. A mineral-supplemented diet increases cortical and trabecular bone mass with high-speed exercise.

## INTRODUCTION

Weight-bearing exercise increases bone mass, structural strength, and tissue quality, making bone better able to resist fracture [1]. Exercise can have long-term benefits to bone health [2–8], making it beneficial to maximize bone mass accumulation early in life. Greater bone mass early in life may be needed to maintain a higher level of bone mass in old age when bone mass declines while weight-bearing exercise becomes more difficult to perform [9–11].

Bone responds to loading from exercise by increasing bone mass and bone strength to accommodate greater loads on bones and prevent damage from future exercise loads [12]. Increasing loading on bones during exercise can increase bone mass since increasing magnitude of strain from loading increases bone cross-sectional area [13] and bone formation rate [14,15]. High-intensity exercise using increased treadmill speed may similarly be able to change loading on the bone by increasing loading frequency. Loading frequency is directly related to bone formation rate, and loads of greater strain magnitude are more effective towards increasing bone formation rate when loads are applied at a higher loading frequency [16].

Increasing bone formation rate may allow bone to reach greater peak bone mass or to achieve peak bone mass in a shorter time, allowing for a more rapid increase in resistance to fracture. This increased bone formation rate could require an increased dietary mineral supply to simultaneously maintain increases in bone mass, bone tissue quality and formation rate with exercise. Since exercise increases demand for dietary minerals [17], high-intensity exercise may cause an even greater need for minerals that normal dietary amounts cannot provide. Combining high-intensity exercise with a calcium- and phosphorus-supplemented diet may be able to maximize bone mass by further increasing bone mass over standard, lower-intensity exercise. Under standard exercise conditions (running at a speed similar to jogging), rodents exercised for 6-12 weeks have increased cortical bone mineral content, area, yield force, and ultimate force [8,18–21]. In young adult mice, combining exercise with a mineral-supplemented diet increases cortical tissue mineral content (TMC) and area compared to exercise with a standard diet after only 3 weeks of exercise [22]. It was hypothesized that combining a mineral-supplemented diet with high-speed treadmill exercise would lead to greater cortical TMC and area than high-speed exercise on a standard diet after 4 weeks of exercise in young adult mice.

## METHODS

### Animals and Treatments

All animal protocols were approved by the University of Michigan University Committee on Use and Care of Animals. One hundred thirty-eight male C57BL/6 mice, 26.3 ± 2.8 g mean body weight, were purchased from Charles River Laboratories (Wilmington, MA) at 13 weeks of age and placed in single housing to prevent fighting. The mice were started on the control diet and were given 2 weeks to acclimate. Starting on experiment day 1, at 15 weeks of age, mice were randomly assigned to one of 7 groups – a baseline group sacrificed on day 1 (B), a non-exercise group fed the control diet (C), a non-exercise group fed the supplemented diet (D), a low-speed exercise group fed the control diet (CE), a low-speed exercise group fed the supplemented diet (DE), a high-speed exercise group fed the control diet (CE+), and a high-speed exercise group fed the supplemented diet (DE+). Mice were divided into groups of equal mean body weight and baseline serum Ca concentration. Baseline serum Ca was measured 5 days before experiment day 1. After 30 days of treatments, all mice from the experimental groups were sacrificed at 19 weeks of age, and left tibiae were harvested for analysis.

### Diets and Exercise Program

The control diet consisted of an AIN-93G diet (TestDiet®, Richmond, IN) modified by adding dicalcium phosphate to contain 0.5% Ca and 0.5% P. The supplemented diet was modified by adding dicalcium phosphate and calcium carbonate to contain 5% Ca and 1% P. Ca, P, and Ca:P ratio were all increased to increase serum Ca by increasing intestinal Ca absorption [23–25]. The control diet contained 3.90 kcal/g with an energy distribution of 65.0% carbohydrates, 16.3% fat, and 18.7% protein while the supplemented diet had 3.39 kcal/g with the same energy distribution. All other nutrients were equivalent between the two diets. The low-speed exercise program consisted of running on a 5° incline treadmill at 12 m/min, 20 min/day for 29 consecutive days. Mice were gradually increased to a maximum speed of 12 m/min in the first 3 days of exercise. The high-speed exercise program consisted of running on a 5° incline treadmill at 20 m/min, 20 min/day. Mice were gradually increased to a maximum speed of 20 m/min in the first 8 days of exercise. Video analysis of mice running gave an estimated average frequency of 3.4 steps/second in mice running 12 m/min and 4.2 steps/second in mice running 20 m/min. Thus, the high-speed exercise had greater loading frequency, and both exercise programs offered a sufficient number of load cycles/day to reach loss of mechanosensitivity such that any further increase in duration would likely not affect bone [26].

### Cortical Geometry and Trabecular Architecture Measurements

Whole tibiae were embedded in 1% agarose, placed in a 19 mm diameter tube, and scanned using a micro-CT specimen scanner (μCT100 Scanco Medical, Bassersdorf, Switzerland) with a voxel size of 12 μm (70 kVp, 114 μA, 0.5 mm AL filter, and integration time 500 ms). Scans were analyzed with Scanco IPL software. A 180-μm thick transverse section from a standard site located 21.7% of the distance from the tibia-fibula junction to the proximal end of the tibia was chosen for measurement of cortical geometry metrics - TMC, volumetric tissue mineral density (TMD), cross-sectional area, and moment of inertia about the anterior-posterior axis. This section is located approximately at the center of the mechanical testing region. Geometry metrics were calculated using a fixed global threshold of 26% (260 on a grayscale of 0–1000) to separate bone from non-bone. Another 180-μm thick transverse section at the fracture site was analyzed for cortical geometry measurements used in calculations of tissue-level mechanical properties (moment of inertia, distance from neutral axis). Tibial scans were further analyzed for trabecular bone architecture. Proximal tibial metaphyseal sections immediately below the growth plate of 480 μm thick were analyzed in Scanco IPL software using freehand traced volumes of interest. Architecture metrics measured were bone volume (BV), bone volume fraction (BV/TV), TMD, Tb. N, Tb. Th, and Tb. Sp. These metrics were calculated using a fixed global threshold of 18% (180 on a grayscale of 0–1000) to separate bone from non-bone.

### Mechanical Testing

Structural- and tissue-level mechanical properties were measured in all groups. Structural-level properties (force, deformation, stiffness, work) were measured from a 4-point bending to failure test (3-mm inner and 9-mm outer spans). Tibiae were loaded to failure with the medial side of the mid-diaphysis in tension under displacement control at 0.025 mm/sec at a data sampling rate of 30 Hz. Tissue-level mechanical properties (stress, strain, modulus, toughness) were estimated using beam bending theory with geometric measurements (moment of inertia about anterior-posterior axis, distance from centroid to medial side of the bone) from micro-CT data at the fracture site [27].

### Serum Analysis

Fasting blood samples taken before daily exercise were collected by submandibular vein bleeding. Blood samples were collected at baseline (day −4), after the first day of full speed running for the high-speed exercised mice (day 9) and on the final day (day 30). Serum was isolated by centrifuge. Ca and P concentrations were measured by using the Calcium CPC LiquiColor test kit (Stanbio Laboratory, Boerne, TX) and the Phosphorus Liqui-UV kit (Stanbio Laboratory). ELISAs were used to measure markers of bone formation and resorption – pro-collagen type I amino-terminal peptide (PINP) and carboxy-terminal collagen crosslinks (CTX) (Immunodiagnostic Systems, Inc., Scottsdale, AZ) – on samples from day 9 and day 30. All manufacturers’ kit instructions were followed, including the use of the standards provided for obtaining standard curves.

### Statistical Analysis

Cortical geometry measurements, trabecular architecture measurements, and mechanical properties were tested by Two-way ANOVA with Tukey’s post-hoc tests to determine if the individual effects of diet or exercise were significant (p < 0.05) and if the combined treatments had a significant interactive effect. Student’s t-tests were used to compare baseline to experimental groups.

## RESULTS

### High-Speed Exercise Prevented Weight Gain after Four Weeks

Exercise had a significant main effect that decreased body weight on day 29 (p < 0.05, Two-way ANOVA, Figure 1). Both of the high-speed exercised groups and mice exercised at low-speed while fed the control diet did not gain weight after day 8. Mice exercised at low-speed while fed the supplemented diet and both of the non-exercised mice groups continued gaining weight after day 8 and the body weights at day 29 of mice in these three groups were significantly higher than the body weights of mice in groups that did not gain weight during the same time (p < 0.05, Tukey’s tests).

**Figure 1.**
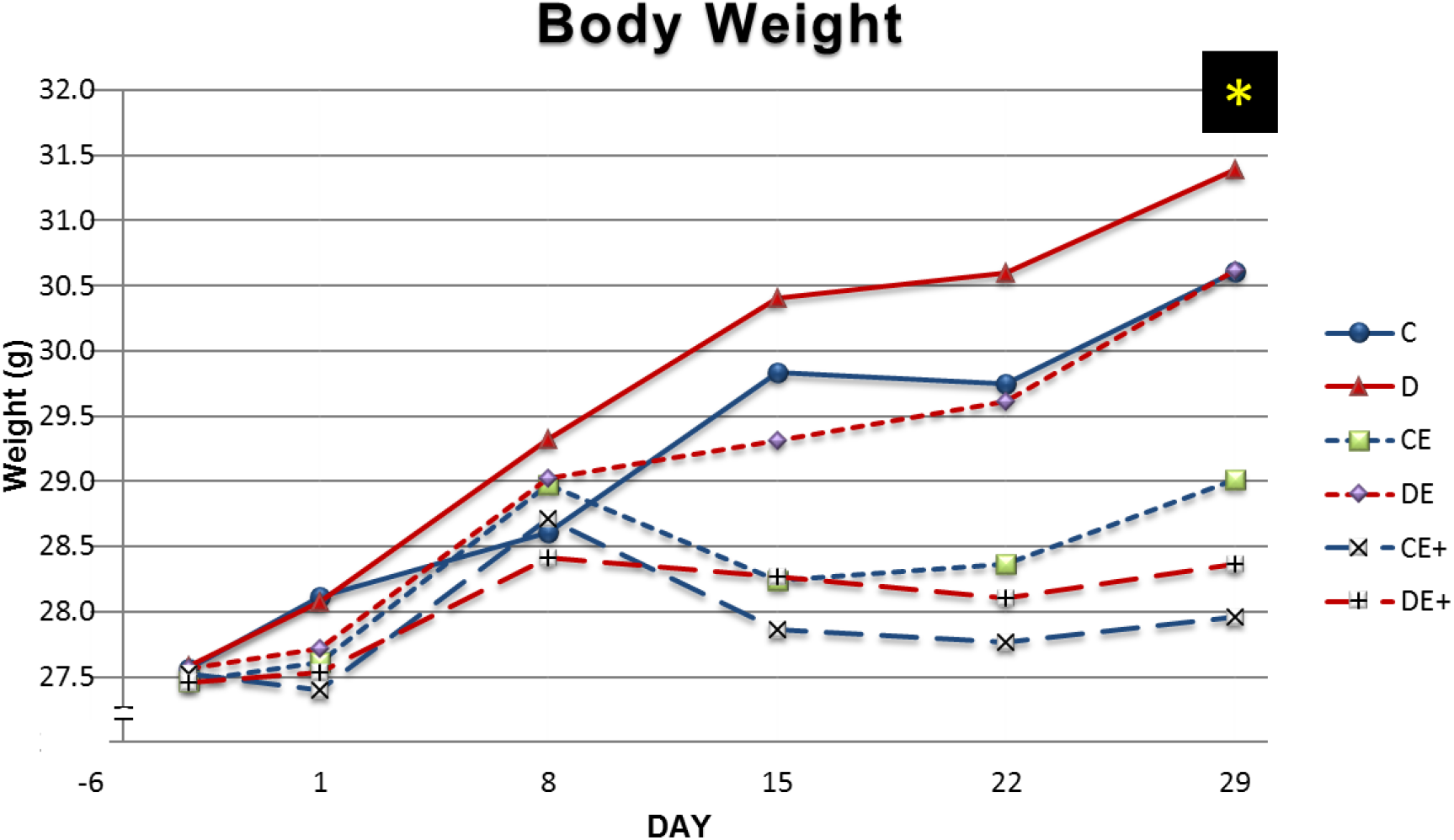
Mouse body weight (mean ± SD). High-speed exercise limited gains in body weight after four weeks. On day 29, both of the high-speed exercised groups and the low-speed exercised group on the control diet had lower body weight than the non-exercised groups and the exercised group on the supplemented diet. *Significant exercise effect on Day 29 (p < 0.05, Two-way ANOVA). C – non-exercised mice fed the control diet, D – non-exercised mice fed the supplemented diet, CE – low-speed exercised mice fed the control diet, DE – low-speed exercised mice fed the supplemented, CE+ – high-speed exercised mice fed the control diet, DE+ – high-speed exercised mice fed the supplemented diet

### The Supplemented Diet Increased Tibial Cortical TMC and Area, and High-Speed Exercise Prevented an Increase in Tibial Length after Four Weeks

Exercise had a significant main effect on tibial length (p < 0.05, Two-way ANOVA, Figure 2). Four weeks of high-speed exercise prevented increases in tibial length for both groups of high-speed exercised mice. Tibial length was significantly higher in low-speed exercised mice on the control diet than high-speed exercised mice on the control diet (p < 0.05, Tukey’s test). All groups except the 2 high-speed exercised mice groups had significantly greater tibial length than baseline (p < 0.05, t-test). There were significant main increasing effects of diet on cortical TMC and area after 4 weeks (p < 0.05, Two-way ANOVA). High-speed exercised mice on the supplemented diet had significantly greater TMC and area than high-speed exercised mice on the control diet (p < 0.05, Tukey’s tests) and baseline mice (p < 0.05, t-test). Tissue mineral density was significantly greater than baseline in all groups except the non-exercised mice on the supplemented diet (p < 0.05, t-test).

**Figure 2.**
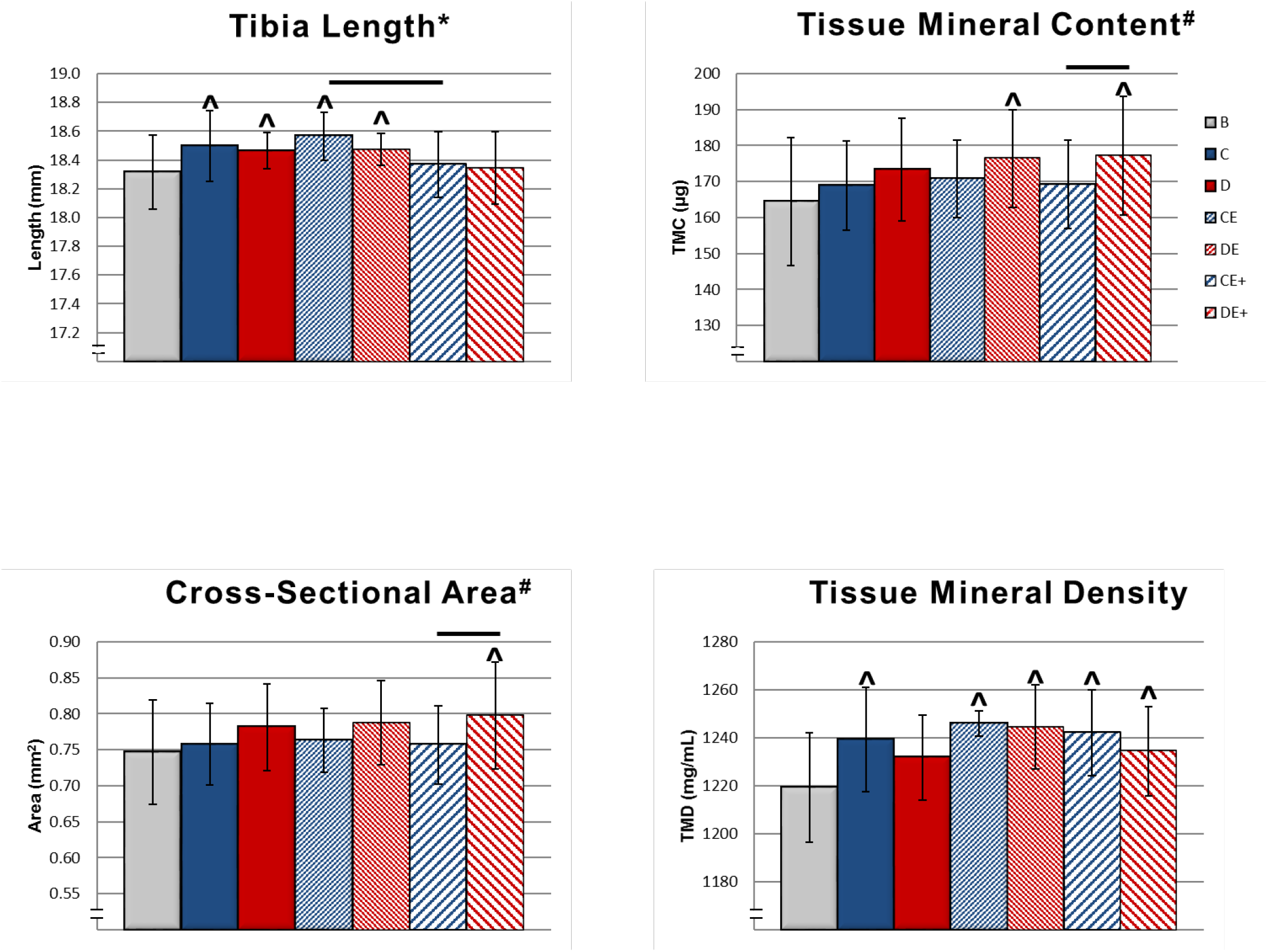
Mouse tibial cortical bone cross-sectional geometric properties and mineralization (mean ± SD). Tibial length increased from baseline for all groups except the high-speed exercised groups. High-speed exercised mice fed the supplemented diet had significantly greater TMC and cross-sectional area compared to high-speed exercised mice fed the control diet. ^#^Significant diet effect (p < 0.05, Two-way ANOVA). *Significant exercise effect (p < 0.05, Two-way ANOVA). ^Significantly different from baseline (p < 0.05, t-test). Horizontal bar represents significant difference between groups (p < 0.05, Tukey’s test). B – baseline mice, C – non-exercised mice fed the control diet, D – non-exercised mice fed the supplemented diet, CE – low-speed exercised mice fed the control diet, DE – low-speed exercised mice fed the supplemented, CE+ – high-speed exercised mice fed the control diet, DE+ – high-speed exercised mice fed the supplemented diet

### The Supplemented Diet Increased Trabecular Bone Volume, and Both Speeds of Exercise Prevented Loss of Trabecular Bone Volume for Mice on the Control Diet after Four Weeks

There were significant main effects of diet that increased trabecular BV, BV/TV, Tb. N, and Tb. Th and decreased Tb. Sp (p < 0.05, Two-way ANOVA, Figure 3). Each group on the supplemented diet had significantly greater trabecular BV, BV/TV, Tb. N, and Tb. Th and significantly lesser Tb. Sp than the control diet group subjected to the same exercise intensity (p < 0.05, Tukey’s tests). Low-speed exercised mice on the control diet had significantly greater BV/TV than non-exercised mice on the control diet. High-speed exercised mice on the control diet had significantly greater Tb. N than non-exercised mice on the control diet. All groups on the supplemented diet had significantly greater BV and BV/TV than baseline (p < 0.05, t-test). The non-exercised mice on the control diet had significantly lower BV and BV/TV than baseline (p < 0.05, t-test). All groups on the control diet had significantly lesser Tb. N and significantly greater Tb. Sp than baseline. Mice from all groups had significantly greater Tb. Th than baseline. Low-speed exercised mice on the supplemented diet and all mice on the control diet had significantly greater TMD than baseline.

**Figure 3.**
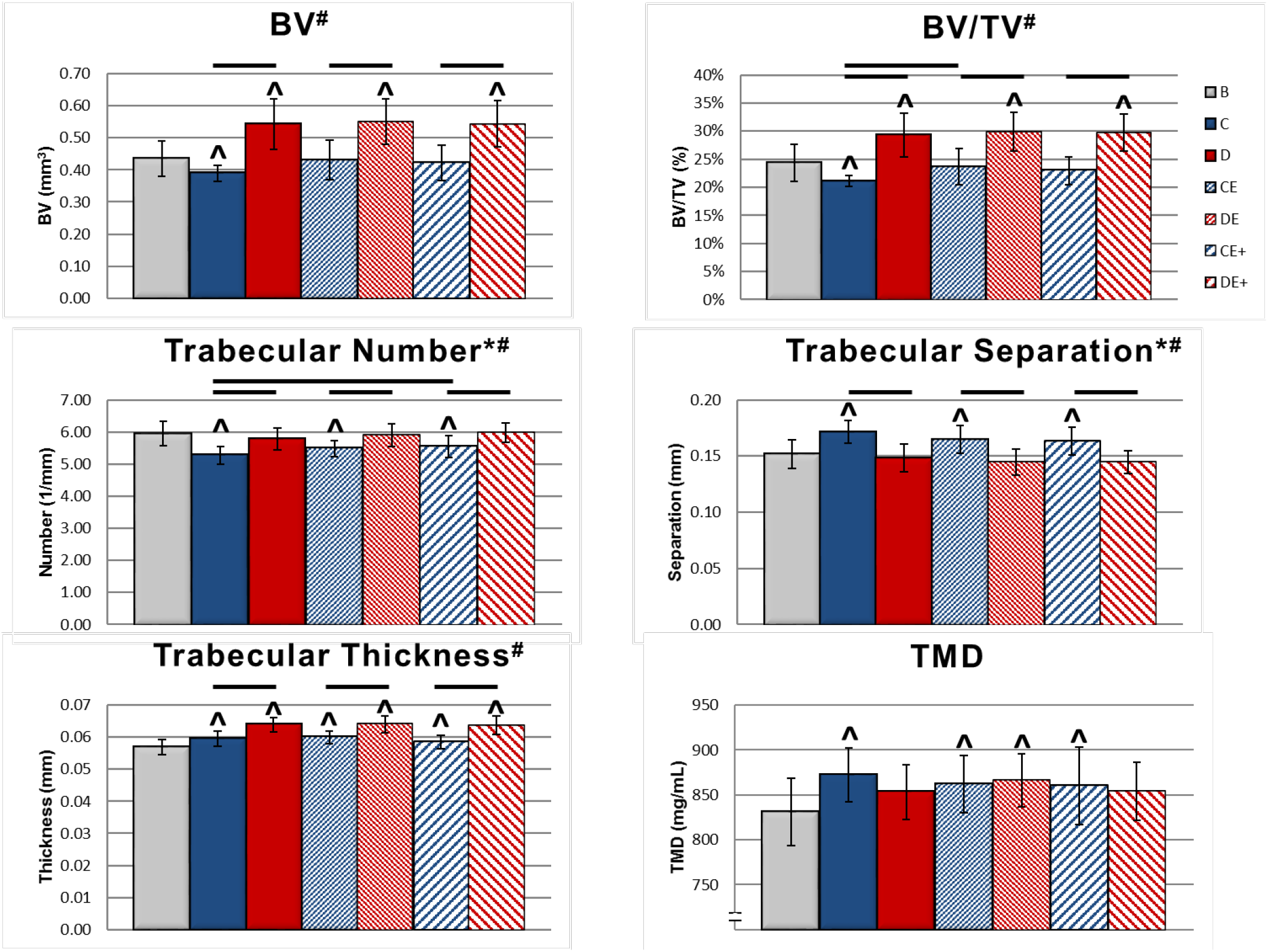
Proximal tibial trabecular architecture (mean ± SD). Diet had a significant main effect on every property except tissue mineral density (TMD), increasing trabecular bone volume (BV), bone volume/total volume (BV/TV), Tb. N, and Tb. Th and decreasing Tb. Sp. For mice on the control diet, both speeds of exercise prevented a significant decrease in BV and BV/TV from baseline. #Significant diet effect (p < 0.05, Two-way ANOVA). *Significant exercise effect (p < 0.05, Two-way ANOVA). ^Significantly different from baseline (p < 0.05, t-test). Horizontal bar represents significant difference between groups (p < 0.05, Tukey’s test). B – baseline mice, C – non-exercised mice fed the control diet, D – non-exercised mice fed the supplemented diet, CE – low-speed exercised mice fed the control diet, DE – low-speed exercised mice fed the supplemented, CE+ – high-speed exercised mice fed the control diet, DE+ – high-speed exercised mice fed the supplemented diet

### The Supplemented Diet Increased Yield Force, and Exercise Had No Effects on Structural-Level Mechanical Properties after Four Weeks

There was a significant main effect of diet that increased yield force after 4 weeks (p < 0.05, Two-way ANOVA, Figure 4). On the supplemented diet, non-exercised mice and low-speed exercised mice had significantly greater yield force and pre-yield work than baseline (p < 0.05, t-test). Low-speed exercised mice on the control diet also had significantly greater yield force than baseline. There were no significant main effects of exercise or significant group differences for any structural-level mechanical property.

**Figure 4.**
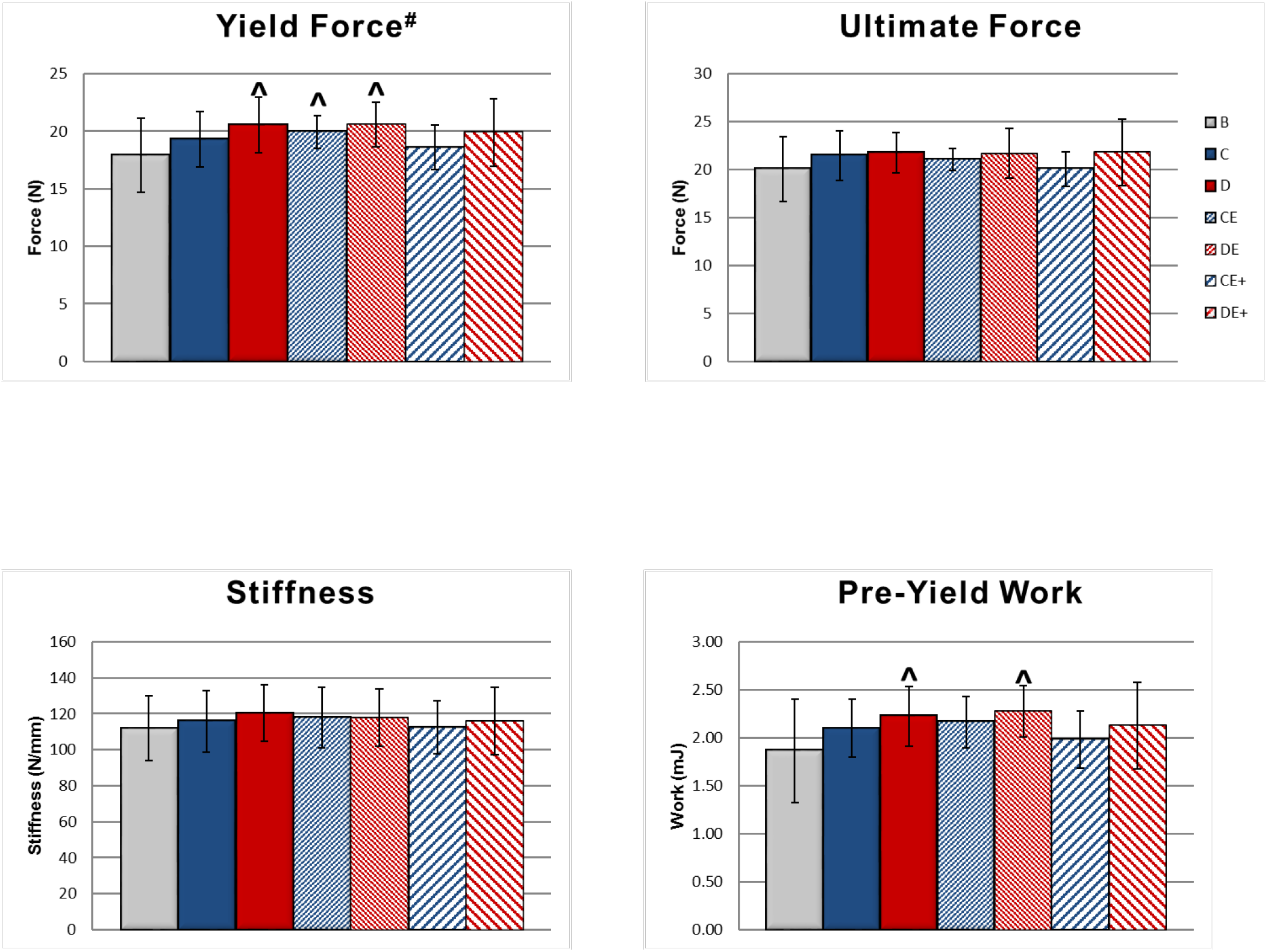
Structural-level tibial mechanical properties (mean ± SD). There was a significant main effect of diet on yield force, and non-exercised mice on the supplemented diet and mice exercised at low intensity on the standard and supplemented diets had significantly greater yield force and pre-yield work than baseline. #Significant diet effect (p < 0.05, Two-way ANOVA). ^Significantly different from baseline (p < 0.05, t-test). B – baseline mice, C – non-exercised mice fed the control diet, D – non-exercised mice fed the supplemented diet, CE – low-speed exercised mice fed the control diet, DE – low-speed exercised mice fed the supplemented, CE+ – high-speed exercised mice fed the control diet, DE+ – high-speed exercised mice fed the supplemented diet

### High-Speed Exercise Decreased Ultimate Stress, and the Supplemented Diet Had No Effects on Tissue-Level Mechanical Properties after Four Weeks

Exercise had a significant main effect that decreased ultimate stress, and pooled ultimate stress data from high-speed exercised mice was significantly lower than from non-exercised mice (p < 0.05, Two-way ANOVA, Figure 5). Ultimate stress was significantly lower in high-speed exercised mice on the supplemented diet than non-exercised mice on the supplemented diet (p < 0.05, Tukey’s test). The non-exercised mice on the supplemented diet had significantly greater yield stress, ultimate stress, and Young’s modulus than baseline (p < 0.05, t-test). There were no significant main effects of diet on tissue-level mechanical properties.

**Figure 5.**
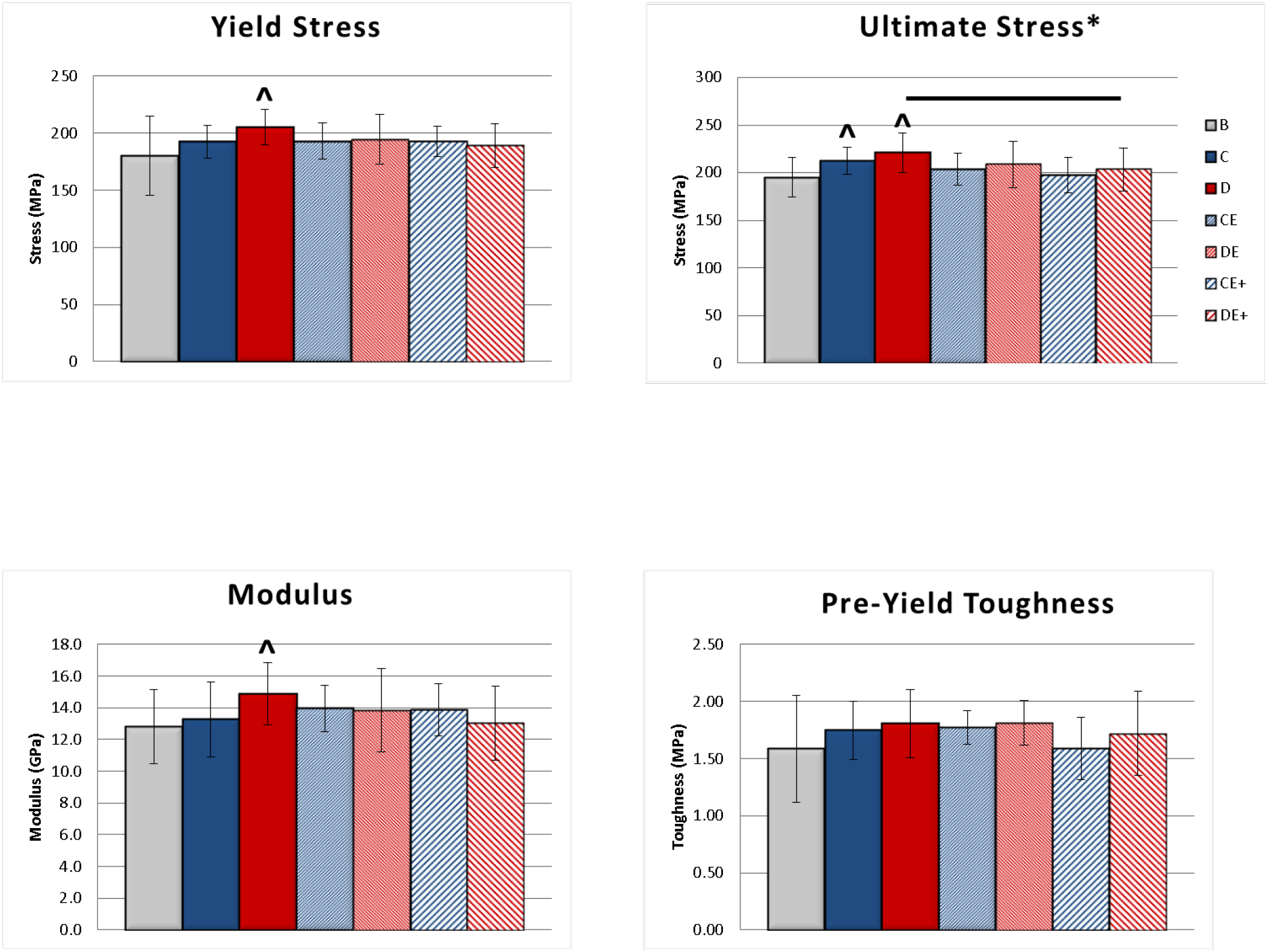
Tissue-level tibial mechanical properties (mean ± SD). Yield stress, ultimate stress, and modulus were all significantly greater than baseline in non-exercised mice on the supplemented diet. Exercise had a significant decreasing main effect on ultimate stress, and high-speed exercised mice on the supplemented diet had significantly lower ultimate stress than non-exercised mice on the supplemented diet. *Significant exercise effect (p < 0.05, Two-way ANOVA). ^Significantly different from baseline (p < 0.05, t-test). Horizontal bar represents significant difference between groups (p < 0.05, Tukey’s test). B – baseline mice, C – non-exercised mice fed the control diet, D – non-exercised mice fed the supplemented diet, CE – low-speed exercised mice fed the control diet, DE – low-speed exercised mice fed the supplemented, CE+ – high-speed exercised mice fed the control diet, DE+ – high-speed exercised mice fed the supplemented diet

### The Supplemented Diet Increased Serum Ca on Day 9 and Day 30 While Exercise Increased Serum Ca only on Day 30

Diet had significant main effects that increased serum Ca on day 9 and day 30, relative to the non-exercised mice on the control diet (p < 0.05, Two-way ANOVA, Figure 6). Exercise had a significant main effect that increased serum Ca on day 30. High-speed exercised mice had significantly greater serum Ca than non-exercised mice on day 30 (p < 0.05, Tukey’s test). There were no significant differences between groups on the supplemented diet at the same time point. There were significant main effects of diet, exercise, and diet and exercise interaction on serum P on day 30. Both exercised groups on the supplemented diet had significantly lower serum P than the corresponding exercised groups on the control diet and lower serum P than the non-exercised mice on the supplemented diet.

**Figure 6.**
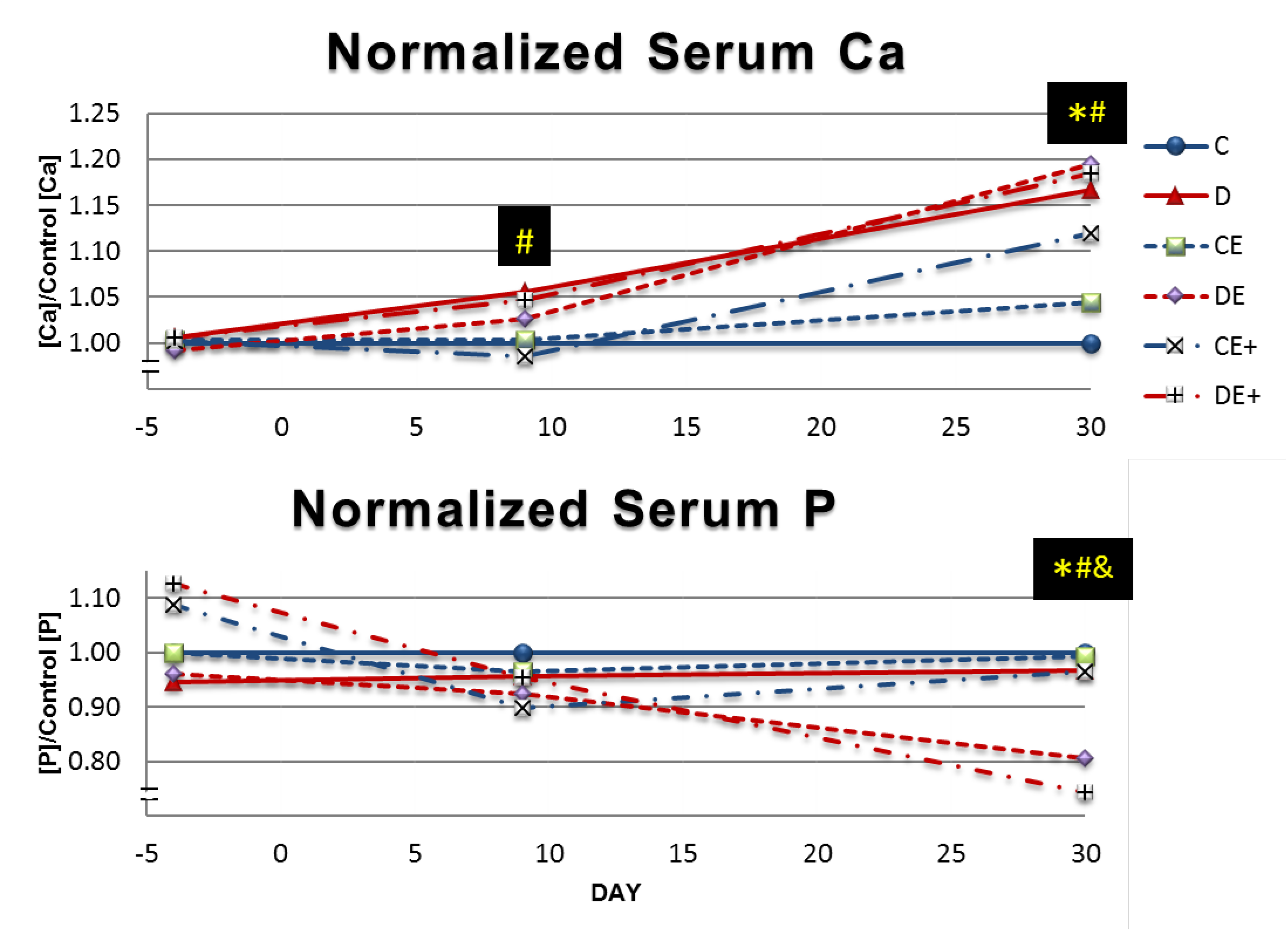
Mean serum [Ca] and [P] on days −4, 9, and 30. Data is normalized to serum concentrations in non-exercised mice on the control diet at the same time point. The supplemented diet significantly increased serum Ca on days 9 and 30. Exercise only significantly increased serum Ca on day 30. Serum P was significantly decreased in exercised mice on the supplemented diet only after day 30. #Significant diet effect on that day (p < 0.05, Two-way ANOVA). *Significant exercise effect on that day (p < 0.05, Two-way ANOVA). &Significant diet and exercise interaction on that day (p < 0.05, Two-way ANOVA). C – non-exercised mice fed the control diet, D – non-exercised mice fed the supplemented diet, CE – low-speed exercised mice fed the control diet, DE – low-speed exercised mice fed the supplemented, CE+ – high-speed exercised mice fed the control diet, DE+ – high-speed exercised mice fed the supplemented diet

### Exercise Decreased PINP/CTX Ratio in Mice on the Control Diet on Day 9, and Mice on the Supplemented Diet on Day 30

Exercise had significant main effects that increased serum CTX and decreased PINP/CTX ratio on day 9 (p < 0.05, Two-way ANOVA, Figure 7). Both groups of exercised mice on the control diet had significantly higher day 9 CTX than the non-exercised mice on the control diet (p < 0.05, Tukey’s test). Non-exercised mice on the control diet had significantly higher day 9 PINP/CTX ratio than non-exercised mice on the supplemented diet and both groups of exercised mice on the control diet. Exercise had significant main effects on serum CTX, serum PINP, and PINP/CTX ratio on day 30. Diet had a significant main effect that increased serum CTX on day 30. There was also a significant diet and exercise interaction on the day 30 PINP/CTX ratio. In high-speed exercised mice on the supplemented diet, day 30 serum CTX was significantly higher, and day 30 CTX/PINP ratio was significantly lower compared to high-speed exercised mice on the control diet. Both day 30 serum CTX and PINP were significantly lower for high-speed exercised mice on the control diet than they were for that group on day 9 (p < 0.05, t-test). Day 30 PINP/CTX ratio was significantly lower for non-exercised mice on the control diet than it was for that group on day 9.

**Figure 7.**
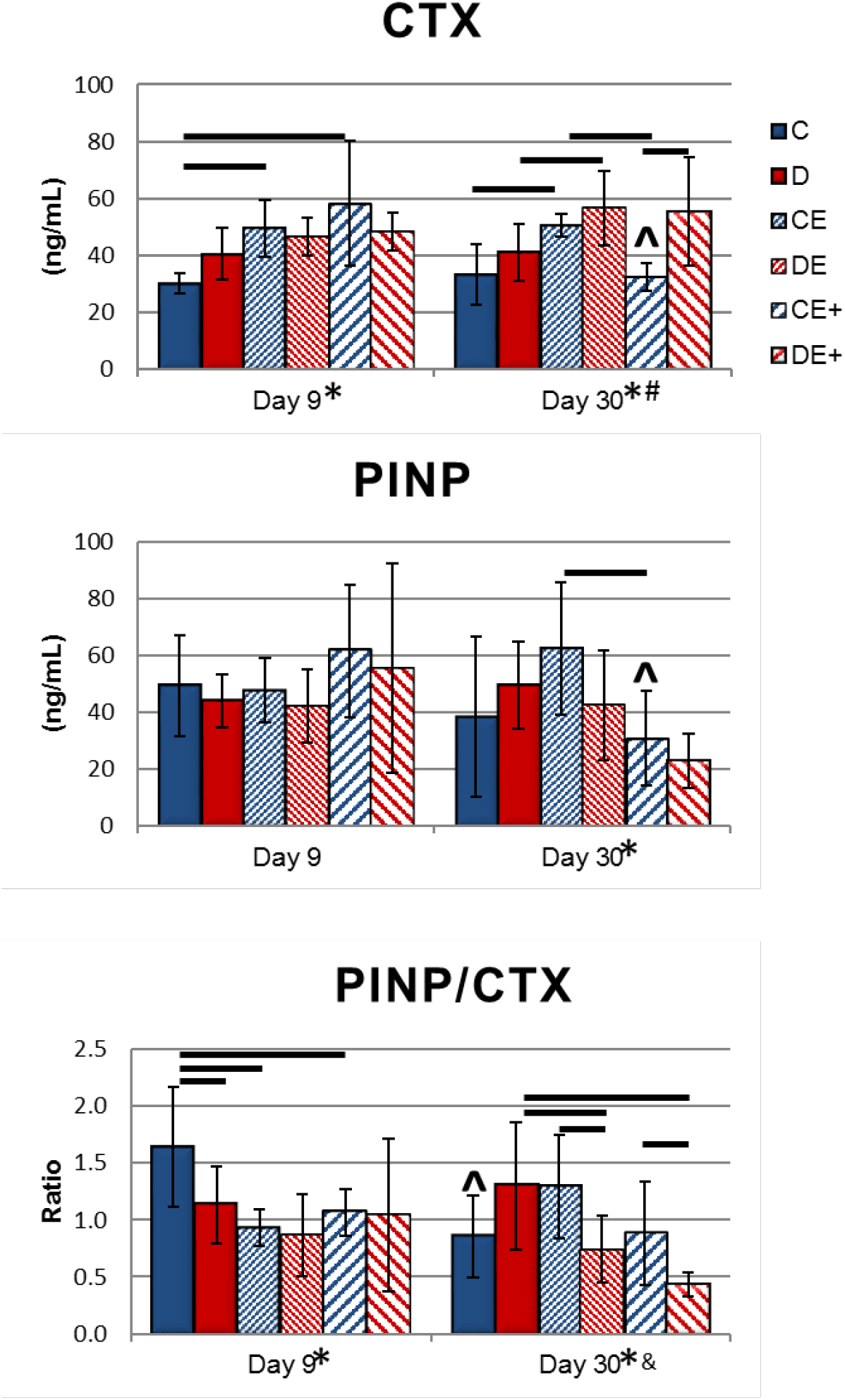
Serum CTX and PINP (mean ± SD) on day 9 and day 30. Exercise significantly increased CTX on day 9 and day 30, regardless of treadmill speed. High-speed exercise significantly decreased PINP on day 30, compared to low-speed exercise. For mice on the control diet, exercise significantly decreased PINP/CTX on day 9, causing a less formation-favored state of bone metabolism. Similarly, for mice on the supplemented diet, exercise significantly decreased PINP/CTX on day 30. #Significant diet effect (p < 0.05, Two-way ANOVA). *Significant exercise effect (p < 0.05, Two-way ANOVA). ^Significant difference between day 9 and day 30 measurement (p < 0.05, t-test). Horizontal bar represents significant difference between groups (p < 0.05, Tukey’s test). &Significant diet and exercise interaction on that day (p < 0.05, Two-way ANOVA).

## DISCUSSION

Four weeks of high-speed exercise prevented longitudinal tibial growth and increases in yield force from baseline (Figure 2, Figure 4). The data suggests that under the higher frequency loading, longitudinal bone growth may have been limited in favor of increasing cortical bone mass and mineralization. When the supplemented diet was added to high-speed exercise, bone may have prioritized increasing cortical bone TMC and area over restoring longitudinal growth. This prioritization may be a sign that the dietary mineral supply was suboptimal for maximizing benefits from exercise.

While for cortical bone there were no main effects of exercise on morphology, mineralization or mechanical properties, and few significant group differences between mice on different diets, trabecular bone was impacted more by the diet and exercise. The supplemented diet increased trabecular bone volume, independent of exercise (Figure 3). Exercise was more effective in mice on the control diet as both speeds of exercise increased BV and BV/TV relative to non-exercised mice. The speed of exercise did not affect trabecular bone properties. The different effects of diet and exercise on trabecular bone volume suggest that while BV and BV/TV are decreasing with age from 15 to 19 weeks, exercise without dietary intervention prevents this decline. Increasing dietary mineral supply increases bone volume, regardless of exercise.

Since combining exercise with the supplemented diet offered no further increases in trabecular bone volume over the supplemented diet alone, exercise may only be affecting trabecular bone when there is a smaller supply of dietary minerals. Mice on the supplemented diet had significantly greater serum Ca than non-exercised mice on the control diet on day 9 and day 30, while exercised mice on the control diet only had significantly greater serum Ca on day 30 (Figure 6). The greater serum supply of Ca from the supplemented diet may have led to greater increases in bone volume at earlier time points, which ultimately led to greater BV and BV/TV after 4 weeks. Exercise alone increased serum Ca only after day 9, and this increased mineral supply may be what prevented the decrease in BV and BV/TV from baseline in exercised mice on the control diet. There could be some peak bone volume achieved such that combining exercise with the supplemented diet did not increase BV beyond what was achieved with the supplemented diet alone. If that is the case, it may be possible that extending the duration of the exercise program could lead to exercised mice on the control diet also reaching some peak bone volume.

Exercise may have only affected trabecular bone because changes in the response of bone to stimuli can be site specific [28]. The increased BV/TV, but not cortical area, TMC or TMD, in low-speed exercised mice on the control diet compared to non-exercised mice on the control diet may be a sign that this short-term exercise program was more effective on trabecular bone. Longer-term exercise for 8 weeks increases cortical TMC and area [22] and may also have increased cortical TMC and area here if exercise had been continued for 4 more weeks. Changes in trabecular, but not cortical bone after a short exercise program could occur because trabecular bone metabolism can be more rapid than cortical bone metabolism.

There were few significant main effects of exercise or diet on structural-level and tissue-level mechanical properties as the supplemented diet increased only yield force, while exercise decreased ultimate stress (Figure 4, Figure 5). This lack of effect on mechanical properties is similar to what was seen after 3 weeks of exercise and supplemented diet treatments [22,29]. After a short duration of an exercise program, bone appears to be in the process of changing tissue composition and increasing bone mass to achieve the changes seen after longer exercise programs of 6-12 weeks. Studying the effects of exercise after 4 weeks may not be adequate for understanding the full impact of exercise on mechanical properties. Also, testing failure strength of bone may not be the best evaluation of short-term exercise effects on bone strength. Fatigue loading bones to determine fatigue life, measure buildup of microdamage, or measure resistance to loss of stiffness with fatigue may be better evaluations of the ability of short-term exercise to increase tissue quality.

There was no significant effect of exercise intensity on any measurements of cortical or trabecular bone, or mechanical properties. Increasing treadmill speed may not be impactful on bone if it does not lead to greater peak loads on bone, which may have occurred here. High-speed exercise did lead to greater average loading frequency, which increases bone formation rate [16]. Bone formation rate was not directly measured, but the bone metabolism markers CTX, PINP, and PINP/CTX can undergo changes that are indicative of increased bone formation [22,30,31]. High-speed exercised mice on the control diet had significantly lower CTX and PINP on day 30 than on day 9 and lower day 30 CTX and PINP compared to low-speed exercised mice on the control diet (Figure 7). High-speed exercise also led to significantly greater serum Ca on day 30 and significantly lower tibia length for mice on the control diet (Figure 6). For mice on the control diet, this high-speed exercise program appears to be lowering bone turnover and preventing longitudinal growth, the opposite of what happens with longer-term exercise [21]. The supplemented diet appears to be preventing effects of high-speed exercise on tibial length and bone metabolism markers as there were no significant differences in any properties measured between the low-speed and high-speed exercised groups on the supplemented diet. More analysis of tissue composition and tissue quality is needed to determine the full extent of effects of this high-speed exercise regimen and how dietary mineral supply interacts with exercise speed.

Both speeds of exercise significantly decreased PINP/CTX ratio for mice on the control diet on day 9, leading to a less formation-favored state of bone metabolism (Figure 7). Similar to exercise significantly decreasing PINP after one day of exercise [22], lower PINP/CTX on day 9 suggests exercise may be limiting bone growth for over a week. These are transient decreases as there are no significant differences in cortical bone mass, and trabecular bone volume is significantly greater in exercised mice after 4 weeks. Exercise did not affect bone metabolism markers for mice on the supplemented diet on day 9. This difference in effects on bone metabolism may be caused by the elevated serum Ca on day 9 seen in all mice on the supplemented diet and may be partially responsible for different effects of exercise on trabecular bone volume, depending on diet.

The opposite effect of diet occurred on day 30 as exercise significantly decreased the PINP/CTX ratio only for mice on the supplemented diet. One possible explanation is that exercised mice on the supplemented diet may be achieving the same peak bone mass as non-exercised mice on the supplemented diet, but at a faster rate. There would likely be some time point where bone metabolism has slowed down for exercised mice on the supplemented diet as was seen on day 30 in this study. Similarly, for non-exercised mice on the control diet, PINP/CTX is significantly lower on day 30 than on day 9, suggesting a decline in bone metabolism rate. Peak bone mass is expected to be lowest in mice in this group, and this shift in bone metabolism may be a sign that these mice are also at or near peak bone mass. As there are many significant effects of exercise and diet on CTX, PINP, and CTX/PINP ratio on day 30, extending the duration of the exercise and diet treatments would likely lead to further changes in bone mass between the groups. Continuing these treatments until bone metabolism has reached some steady state would allow for the best evaluation of long-term effects of exercise intensity.

Both high-speed and low-speed exercised mice on the supplemented diet had significantly lower day 30 serum P than non-exercised mice and exercised mice on the control diet (Figure 6). It may be possible that serum P was being depleted to use for increasing bone mineralization as both groups with lower serum P were the only groups with significantly higher cortical TMC than baseline (Figure 2). Rodents subjected to treadmill exercise can have increased demand for P as well as Ca [32]. Although the supplemented diet has twice the P as the control diet, this still may not be sufficient for the increased mineral demands from exercise at this time point. Exercise did not decrease serum P for mice on the control diet, but this may be due to the difference in Ca in the diets as the control diet had equal amounts of Ca and P, while the supplemented diet had 5 times more Ca than P. The Ca:P ratio in the supplemented diet may provide an adequate amount of Ca, but not P. Further analysis of tissue composition of these bones may be needed to determine if there were any differences in bone Ca and P content from the diets or exercise treatments. Additionally, mice were not assigned to groups of equivalent baseline serum P. Initial differences in serum P may have been a factor in the significant group differences seen on day 30, though only the exercised mice on the supplemented diet had a decrease in serum P from day 9 to day 30.

One significant main effect of exercise that did not translate into differences in cortical or trabecular bone properties was the lower body weight for all high-speed exercised mice and for low-speed exercised mice on the control diet (Figure 1). Although mice in these groups had lower body weight, they did not have lower tibial bone mass or mechanical strength, as might be expected [33,34]. When bone mass was normalized by weight, the high-speed exercised mice on the supplemented diet had significantly higher tibial length, cortical TMC, area and TMD than all other mice on the supplemented diet (data not shown). Thus, even though exercise did not directly affect most measurements of bone mass or mechanical strength for mice on the supplemented diet, the high-speed exercise allowed for these mice to reach the same bone mass and strength while at a lower body weight. This high-speed exercise regimen may be offering additional health benefits not measured, such as preventing increases in fat mass, increasing lean body mass, and improving cardiovascular health, at no expense to bone health.

After 4 weeks of high-speed treadmill exercise, exercised mice had lower body weight, and the exercise prevented an increase in tibial length, regardless of dietary mineral supply. These changes in growth did not come at the expense of tibial cortical bone mass, trabecular bone volume, or mechanical properties. High-speed exercised mice on the supplemented diet had the greatest cortical TMC and area and trabecular BV and BV/TV after 4 weeks. Combining a high-intensity exercise program with a mineral-supplemented diet may best enable peak bone mass to be reached and prevent fractures.

## Acknowledgments

This work was supported by the NIH award R01-AR052010 and the National GEM Consortium Ph.D. Engineering Fellowship. Thank you to Michelle Lynch and the University of Michigan School of Dentistry MicroCT core, funded in part by NIH/NCRR S10RR026475-01. I would also like to thank Erin McNerny and Joe Gardinier for their guidance and assistance.

